# Task-evoked pulse wave amplitude tracks cognitive load

**DOI:** 10.1101/2023.06.16.545290

**Authors:** Yuri G. Pavlov, Anastasia S. Gashkova, Dauren Kasanov, Alexandra I. Kosachenko, Alexander I. Kotyusov, Boris Kotchoubey

## Abstract

Cognitive load is a crucial factor in mentally demanding activities and holds significance across various research fields. This study aimed to investigate the effectiveness of pulse wave amplitude (PWA) as a measure for tracking cognitive load and associated mental effort in comparison to heart rate (HR) during a digit span task. The data from 78 participants were included in the analyses. Participants performed a memory task in which they were asked to memorize sequences of 5, 9, or 13 digits, and a control task where they passively listened to the sequences. PWA and HR were quantified from photoplethysmography (PPG) and electrocardiography (ECG), respectively. Pupil dilation was also assessed as a measure of cognitive load. We found that PWA showed a strong suppression with increasing memory load, indicating sensitivity to cognitive load. In contrast, HR did not show significant changes with task difficulty. Moreover, when memory load exceeded the capacity of working memory, a reversal of the PWA pattern was observed, indicating cognitive overload. In this respect, changes in PWA in response to cognitive load correlated with the dynamics of pupil dilation, suggesting a potential shared underlying mechanism. Additionally, both HR and PWA demonstrated a relationship with behavioral performance, with higher task-evoked HR and lower PWA associated with better memory performance. Our findings suggest that PWA is a more sensitive measure than HR for tracking cognitive load and overload. PWA, measured through PPG, holds significant potential for practical applications in assessing cognitive load due to its ease of use and sensitivity to cognitive overload. The findings contribute to the understanding of psychophysiological indicators of cognitive load and offer insights into the use of PWA as a non-invasive measure in various contexts.

## Introduction

Cognitive load refers to the amount of working memory (WM) resources required to process information or perform tasks. In a typical WM task, such as the digit span task, the sequential encoding and maintenance of a string of digits incrementally increases cognitive load, culminating in overload when WM capacity is surpassed (Jacobs, 1887). While cognitive load is often gauged through self-report, this subjective assessment is prone to distortion, as it is retrospectively completed post-task (Bermúdez & Massin, 2023; Gendolla & Richter, 2010). Nevertheless, the real-time monitoring of cognitive load and corresponding mental effort is potentially achievable via physiological recordings (Granholm et al., 1996; Kahneman & Beatty, 1966; Peavler, 1974). This approach not only provides valuable insights into the underpinnings of cognition but also offers practical benefits in the ongoing assessment of mental states, having potential applications in various fields, from educational settings to workplace productivity.

Mental effort expended during cognitive tasks often results in changes in autonomic nervous system (ANS) activity. The allocation of resources during demanding task performance is evident through various indicators of cardiovascular activity regulated by the ANS (Gendolla & Wright, 2012). Notable among these indicators are heart rate (HR) (Al abdi et al., 2018; Schapkin et al., 2012; Vincent et al., 1996), pulse wave amplitude (PWA) (Xuan et al., 2020), heart rate variability (HRV) (Mandrick et al., 2016), pre-ejection period and systolic blood pressure (Richter et al., 2008). However, the effectiveness of these indicators in assessing cognitive load and cognitive overload varies based on several factors: the relative contribution of sympathetic and parasympathetic branches of the ANS, the complexity of obtaining the measure, and the effects of individual differences and situational factors.

Cognitively demanding tasks affect cardiovascular indicators mediated by sympathetic rather than parasympathetic influences (Gendolla & Wright, 2012). However, systolic blood pressure serves as an indicator of primarily sympathetic and, less directly, parasympathetic influences (Gendolla & Wright, 2012). Similarly, heart rate (HR) has regulation by sympathetic mechanisms and parasympathetic mechanisms, with parasympathetic dominance observed during minor and moderate efforts (Wright & Kirby, 2001). Thus, recording HR and systolic blood pressure over longer epochs can average sympathetic and parasympathetic effects, which may interact with, or even cancel out, each other.

On the other hand, peripheral vasoconstriction detected via decreased pulse wave amplitude extracted from photoplethysmography (PPG) has primarily sympathetic influence (Babchenko et al., 2001; Shelley, 2007). Other indicators of sympathetic influence are pre-ejection period and some measures of HRV. However, measuring pre-ejection period requires simultaneous recording of ECG and impedance cardiogram or seismocardiogram, and HRV can only be assessed over a long period of stationary activity (at least one minute (Quintana et al., 2016)). To the contrary, PPG is easily accessible (e.g., utilizing a smartphone camera (Matsumura & Yamakoshi, 2013)) and can potentially detect changes in cognitive load over a period of seconds.

Despite its potential, the utilization of pulse wave amplitude in psychological research has been limited. The low adoption of this method may be attributed to individual differences in the absolute values of pulse wave amplitude, which can reduce the effect sizes when comparing different task conditions. Previous studies have observed a decrease in pulse wave amplitude during tasks of increasing difficulty, such as the Sternberg working memory task (Iani et al., 2004), Stroop task (Tulen et al., 1989), and mental arithmetic (Goldstein & Edelberg, 1997). These studies averaged activity over a long period of time related to the task, comparing it with a resting state or tasks of varying difficulty. However, physiological responses may involve alternating periods of increase and decrease in amplitude, which could be obscured when averaged over time. Unfortunately, the long-standing tradition in psychophysiological research that focuses on task-evoked activity changes over time has not been tested in relation to pulse wave amplitude, whereas this method has the potential for enhancing the temporal precision of physiological estimates. Additionally, the method of baseline normalization, particularly when applied to the prestimulus interval in individual trials, can account for individual differences in vascular physiology and situational factors, thereby improving the reliability and validity of PPG measurements in psychological research.

The aim of this study was to explore how cardiovascular responses to cognitive tasks can provide real-time insights into cognitive load. Specifically, we aimed to characterize cardiovascular responses with high precision in time during performance of the digit span task of varying difficulty. We quantified absolute and, to account for individual differences in peripheral physiology, baseline normalized event-related pulse wave amplitude. For comparison, we also report event-related HR extracted from both PPG and ECG sensors. Additionally, we made a comparison with a more established measure of ANS activity, where quantifying task-evoked changes are the norm – pupil dilation (van der Wel & van Steenbergen, 2018).

## Methods

The methods partially overlap with our previously published articles (Kosachenko et al., 2023; Pavlov et al., 2022) where the sample, task, and procedure, as well as the pupillometry data acquisition and analysis are described in greater detail.

### Participants

Eighty-six participants completed the task. The participants had normal or corrected-to-normal vision and reported no history of neurological or mental diseases. All the participants were Russian native speakers. Informed consent was obtained from each participant. The experimental protocol was approved by the Ural Federal University ethics committee. The experiment was conducted in accordance with the Declaration of Helsinki.

Eight participants were excluded because of technical issues or insufficient data quality in one or more conditions (including resting state). Thus, data of 78 participants (69 females, age: 20.5 ± 3.9 years) entered the analyses.

### Task and procedure

Before the WM task, we recorded 4 minutes of eyes-closed resting state ECG and PPG. After that, the participants were given instructions and proceeded with the WM task (see Figure 1; for a more detailed description of the task see Kosachenko et al. (2023)). In this task, participants were either required to memorize sequences of 5, 9, or 13 digits (memory condition) or to listen to them (control condition) without trying to memorize them. Afterward, they were asked to recite each digit in the order in which it was presented (i.e., serial recall). There were 54 control trials and 108 memory trials in 9 blocks separated by self-paced breaks. Each block consisted of 3 control (one of each sequence length) followed by 12 memory (4 trials on each sequence length in random order) followed again by 3 control trials. To evaluate behavioral performance, we implemented partial scoring, which gives points for each digit recalled in its correct position in the sequence. To assess subjective workload, participants completed the NASA-TLX questionnaire (Hart & Staveland, 1988) after every third block of the task.

**Figure 1.**
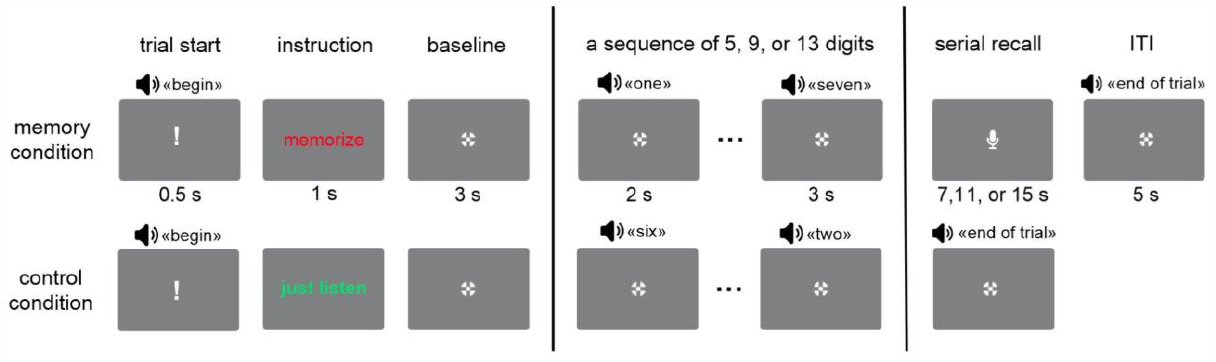
Schematic representation of the English equivalent of the digit span task (stimuli are not to scale). The digits were presented auditorily every 2 s. After the last digit in the sequence, there was 3 s retention interval.

### Cardiovascular measures

Cardiovascular measures (ECG and PPG) were continuously acquired using an actiCHamp amplifier (Brain Products) from the auxiliary (AUX) inputs. The ECG was recorded using one channel with the active electrode on the right wrist, the reference electrode on the left wrist, and the ground electrode on the left inner forearm 3 cm distal from the elbow. Finger photoplethysmography was recorded from the left index finger.

Preprocessing of PPG and ECG included the following steps. First, both PPG and ECG data were filtered between 0.5 and 100 Hz, and a notch filter 50 Hz was applied using *eegfiltnew* function in EEGLAB. Then, the data were downsampled to 250 Hz. Next, HRVTool v.1.07 (Vollmer, 2019) was used to detect R peaks in ECG and identify troughs and peaks in PPG. Each trough and peak was visually inspected and incorrectly extracted values were rejected from subsequent analysis. Inter-beat intervals (IBI) < 0.3 and IBI > 2.4 s, as well as HR and PWA values above and below 3 SDs were also rejected. Pulse wave amplitude was defined as the amplitude difference between consecutive trough and systolic peaks, starting from the trough. After that, the recording was segmented into trials starting from 5.5 s before first digit presentation and ending 12 s after last digit presentation. In the epochs, the instantaneous HR (60/IBI) and PWA series were linearly interpolated in 0.5 s bins to obtain HR and PWA series with equally spaced values. Trials with no valid values were removed from the analysis. Out of 12619 trials, 14 were excluded based on this criterion from the HR (ECG) data analysis, 274 from the HR (PPG), and 324 from the PWA analysis.

For the time-locked analyses, after averaging in trials, 3s time interval before first digit onset was used for baseline normalization (averaged in tasks and sequence lengths separately for each participant). Absolute baseline normalization (subtraction) was used for HR values and relative (percent change) normalization was used for PWA values.

### Statistics

We used a repeated-measures ANOVA for the analysis of the effect of Task (control, memory) and Load (13 levels corresponding to processing of individual digits irrespectively of the sequence length) on HR (extracted from ECG) and PWA. In this analysis, due to the difficulty of considering different levels of Load, results were averaged across all sequence lengths. Another ANOVA comparing baseline normalized and raw data included factors Task and Sequence (3 levels: 5, 9, and 13 digits). Greenhouse-Geisser adjusted degrees of freedom are reported where applicable.

Pearson correlations were calculated to investigate the relationships among behavioral accuracy, NASA-TLX and the physiological variables. cocor package for R (Diedenhofen & Musch, 2015) was used to compare the correlation coefficients between different measures, which is designed for the comparison of overlapping anddependent correlations. We used Hittner, May, and Silver’s modification of Dunn and Clark’s z using a backtransformed average Fisher’s Z procedure to make the comparison (see Diedenhofen & Musch (2015) for references).

The statistical analyses were conducted in R (v. 4.2.2).

## Results

### Performance

The average number of correctly recalled digits in the correct serial position decreased as the sequence length increased. Specifically, the mean (±SD) for 5-digit sequences was 4.62 ± 0.31, for 9-digit sequences it was 4.18 ± 1.18, and for 13-digit sequences it was 3.57 ± 1.13, resulting in an overall mean of 4.12 ± 0.81. Full results of the behavioral data analyses are reported elsewhere (Kosachenko et al., 2023).

### Task and load effects

The HR extracted from PPG and ECG had similar temporal trajectories (Figure 2c,d) and strongly correlated (r(71) = 0.951, p < 0.001). Therefore, we analyzed statistically only HR from ECG to reduce the number of redundant analyses. In the following set of analyses, we used baseline normalized HR and PWA data averaged over all sequences. The differences between the temporal trajectories for the first 5 (ANOVA with three levels of the factor Sequence) and 9 digits (Sequence with two levels) in the sequence was not significant for any comparison. Therefore, we averaged over the sequence length.

**Figure 2.**
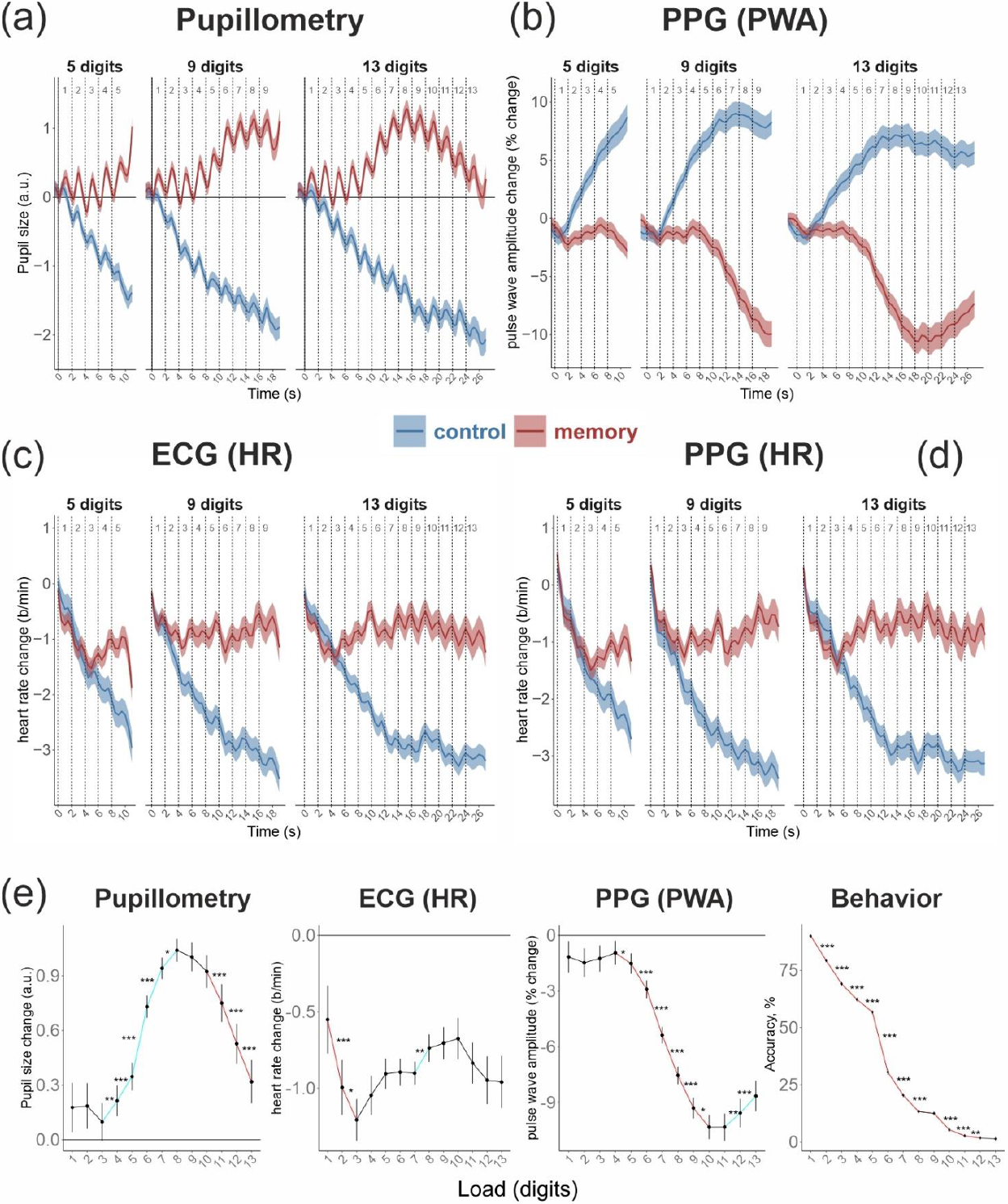
Temporal dynamics of (a) pupil size; (b) pulse wave amplitude (PWA) registered by PPG; (c) heart rate registered by ECG; and (d) heart rate registered by PPG. Each dashed line represents the onset of a digit presentation. Time 0 indicates presentation of the first digit in the sequence. (e) Pupil size, heart rate (from ECG), and PWA in the memory task averaged in the two-second time intervals corresponding to processing of individual digits in the sequence, and behavioral performance in the task averaged over all conditions. Red lines mark a significant decrease and blue lines a significant increase in the parameter compared with the adjacent levels of load. * p<0.05, ** p<0.01, *** p<0.001.

HR was generally higher during the memory task compared to the control task, as indicated by a significant main effect of Task (see Table 1). The effect of Load, however, was only observed in the control task, showing a significant relationship (F(4.02, 309.47) = 61.52, 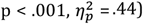). During the memory task, HR exhibited an initial deceleration followed by acceleration, eventually reaching a plateau (Figure 2e). However, Load did not significantly affect HR in the memory task (F(2.21, 170.20) = 1.58, p = .206, 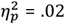). The pairwise comparisons between levels of Load and Tasks are presented in Table S2. The HR effects did not depend on the type of baseline normalization. Table S3 and Figure S1 present the results of percent change normalization applied to interbeat interval data.

**Table 1.**
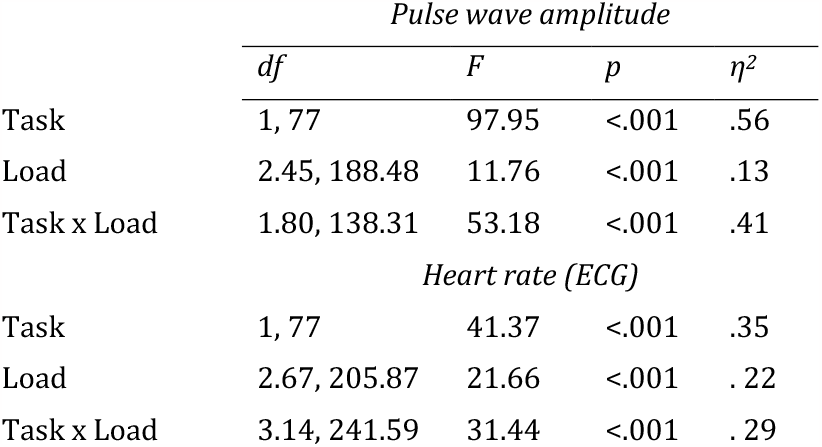
The Task (control vs memory) and Load (13 levels) effects on PWA and HR.

| *Pulse wave amplitude* |  |  |  |  |
| --- | --- | --- | --- | --- |
| | *df* | *F* | *p* | $\eta^2$ |
| Task | 1, 77 | 97.95 | <.001 | .56 |
| Load | 2.45, 188.48 | 11.76 | <.001 | .13 |
| Task x Load | 1.80, 138.31 | 53.18 | <.001 | .41 |
| *Heart rate (ECG)* |  |  |  |  |
| Task | 1, 77 | 41.37 | <.001 | .35 |
| Load | 2.67, 205.87 | 21.66 | <.001 | .22 |
| Task x Load | 3.14, 241.59 | 31.44 | <.001 | .29 |

PWA showed different patterns in the control and memory tasks (Table 1). In the memory task, PWA mostly decreased with increasing load (F(1.75, 134.86) = 35.64, p < .001, 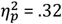; Figure 2b), while in the control task, it mostly increased (F(2.65, 203.99) = 25.57, p < .001, 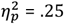; see Table S1 for post hoc tests in both Tasks). As can be seen in Figure 2e, in the memory task, PWA initially dropped after the first digit and then remained relatively stable until the fifth digit, after which there was a significant decrease. The pattern reversed after the 12th and 13th digits in the sequence.

### The effect of baseline normalization

To assess the effect of baseline normalization on the cardiovascular measures, we separately averaged raw and baseline normalized PWA and HR during memorizing or listening to the entire sequence. By averaging over the whole sequence, we were able to compare the results more directly with previous studies, in which large time intervals corresponding to the entire task are typically analyzed.

The most notable difference was observed in the estimation of PWA (Table 2). Interestingly, due to a cross-over Task x Sequence interaction effect in raw data, main effects were not significant (Figure 3, bottom left panel). Specifically, in the case of 5-digit sequences, raw PWA was larger in the memory task compared to the control task (p = 0.039). In the 9-digit sequences it was, in contrast, slightly (not significantly; p = 0.093) larger in the control task, and the same effect was highly significant in the 13-digit sequences (p = 0.002). After baseline normalization, the effect was highly consistent in all conditions, PWA being significantly larger in the control task than in the memory task. Furthermore, normalization visibly reduced the variability of the data, which may have led to the fact that, within the memory task, the difference between the 9 and 13-digit sequences became significant (p = 0.006) while it was not in raw data (p = .231).

**Table 2.**
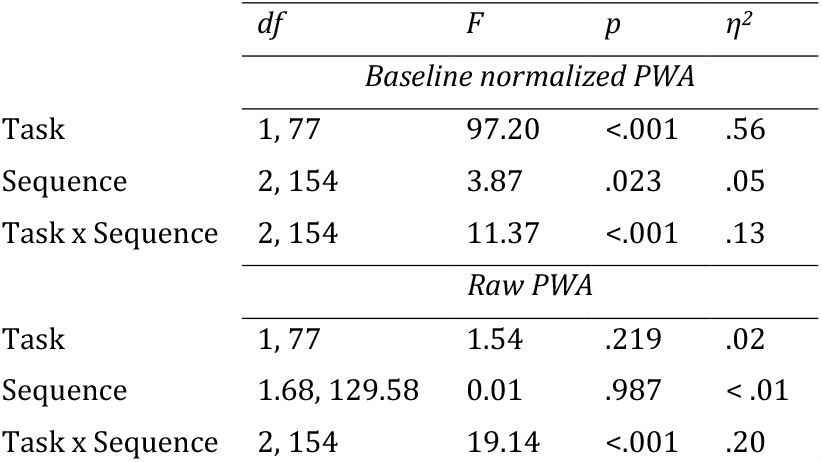
The comparison of baseline normalized and raw PWA values between sequences of 5, 9, and 13 digits in the control and memory tasks.

**Figure 3.**
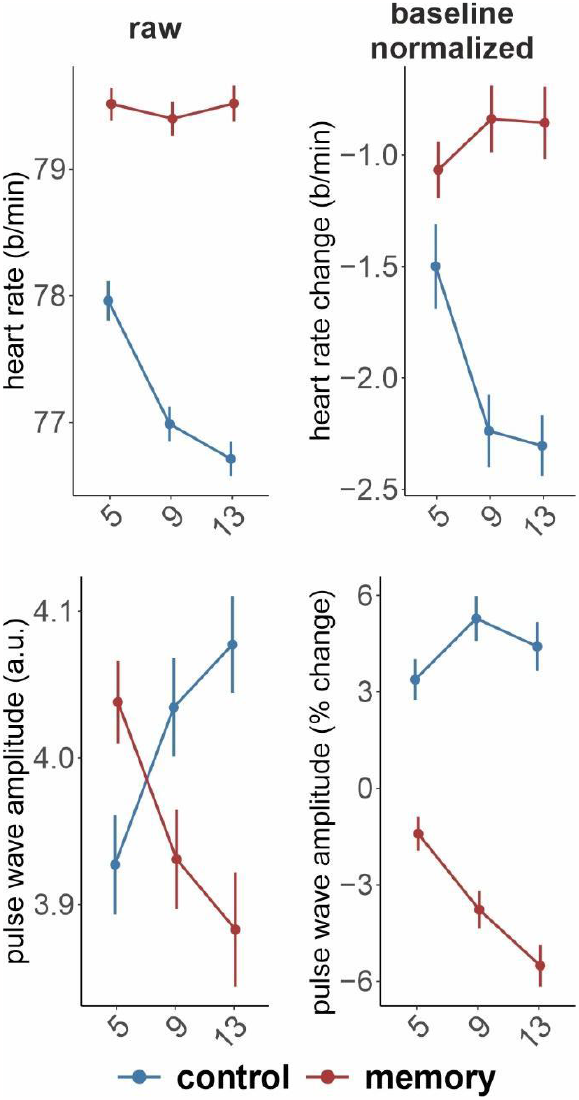
A comparison of baseline normalized and raw HR and PWA in the sequences of 5, 9, or 13 digits.

HR showed a significant decrease in the control task compared to the memory task (Table 3). HR decreased as the sequence length increased in the control task: there was a significant difference in HR between 5-digit sequences and both 9- and 13-digit sequences, as observed in both baseline normalized and raw data (highest p = 0.002). However, in the memory task, neither the baseline normalized nor raw data showed significant pairwise differences in HR between different sequence lengths. Nevertheless, the difference between the tasks remained significant for every sequence length, considering both baseline normalized and raw data.

**Table 3.** The comparison of baseline normalized and raw HR (extracted from ECG) values between sequences of 5, 9, and 13 digits in control and memory tasks.

| | *df* | *F* | *p* | $\eta^2$ |
| --- | --- | --- | --- | --- |
| *Baseline normalized HR* |  |  |  |  |
| Task | 1, 77 | 27.72 | <.001 | .26 |
| Sequence | 2, 154 | 2.68 | .072 | .03 |
| Task x Sequence | 2, 154 | 9.73 | <.001 | .11 |
| *Raw HR* |  |  |  |  |
| Task | 1, 77 | 100.22 | <.001 | .57 |
| Sequence | 1.77, 136.53 | 25.54 | <.001 | .25 |
| Task x Sequence | 1.80, 138.30 | 26.90 | <.001 | .26 |

To investigate the changes in raw PWA and HR compared to the resting state, we conducted an ANOVA with three levels of Task factor: memory, control, and rest. As expected, the effects of Task were significant on both PWA (*F*(1.05, 80.82) = 15.83, *p* < .001, 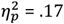) and HR (*F*(1.19, 91.62) = 11.32, *p* < .001, 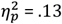).

PWA during rest was significantly larger compared to both memory (t(77) = 4.152, p < 0.001, d = 0.473) and control tasks (t(77) = 3.844, p < 0.001, d = 0.438), while the difference between memory and control tasks was marginally significant (t(77) = 2.007, p = 0.048, d = 0.229). The corresponding means (± standard deviations) were 5.14±3.76, 3.93±2.16, 4.03±2.16, for rest, memory, and control, respectively.

Regarding HR, there was a significant difference between the rest and memory tasks (mean+SD; rest: 77.8±11.5, memory: 79.5±10.9 b/min; t(77) = 2.627, p = 0.01, d = 0.29). However, no significant difference was observed between the rest and control tasks (control: 77±10.5 b/min; t(77) = 1.217, p = 0.227, d = 0.139). Additionally, the control task had significantly lower HR compared to the memory task (t(77) = 10.64, p < 0.001, d = 1.212).

### The relationship between physiological responses and memory accuracy

Next, to understand the relationship between cardiovascular reactivity measures and behavioral performance, we conducted a correlational analysis. In this analysis, we used PWA and HR values averaged across listening and memorizing all 13 digits, along with the average number of correctly recalled digits in the correct order. Additionally, to compare with a well-established index of peripheral nervous system activity known to be associated with cognitive load and effort, we incorporated pupillometry data. Detailed pupillometry data analysis can be found elsewhere (Kosachenko et al., 2022). Here, a figure displaying pupil size dynamics is presented alongside the PPG and ECG data (Figure 2a).

The results revealed significant associations between pupil size and both HR and PWA (Figure 4). In the memory task, the correlation between pupil size and PWA displayed a numerically stronger relationship compared to the pupil size-HR correlation, although the difference did not reach statistical significance (z = -1.627, p = 0.052). Moreover, all physiological variables demonstrated correlations with accuracy in the task. Among the physiological variables examined, only PWA significantly correlated with subjective assessment of cognitive load (r(76) = -0.23, p = 0.038) as measured by the NASA-TLX scale (see the extended correlation matrix in Figure S3).

**Figure 4.**
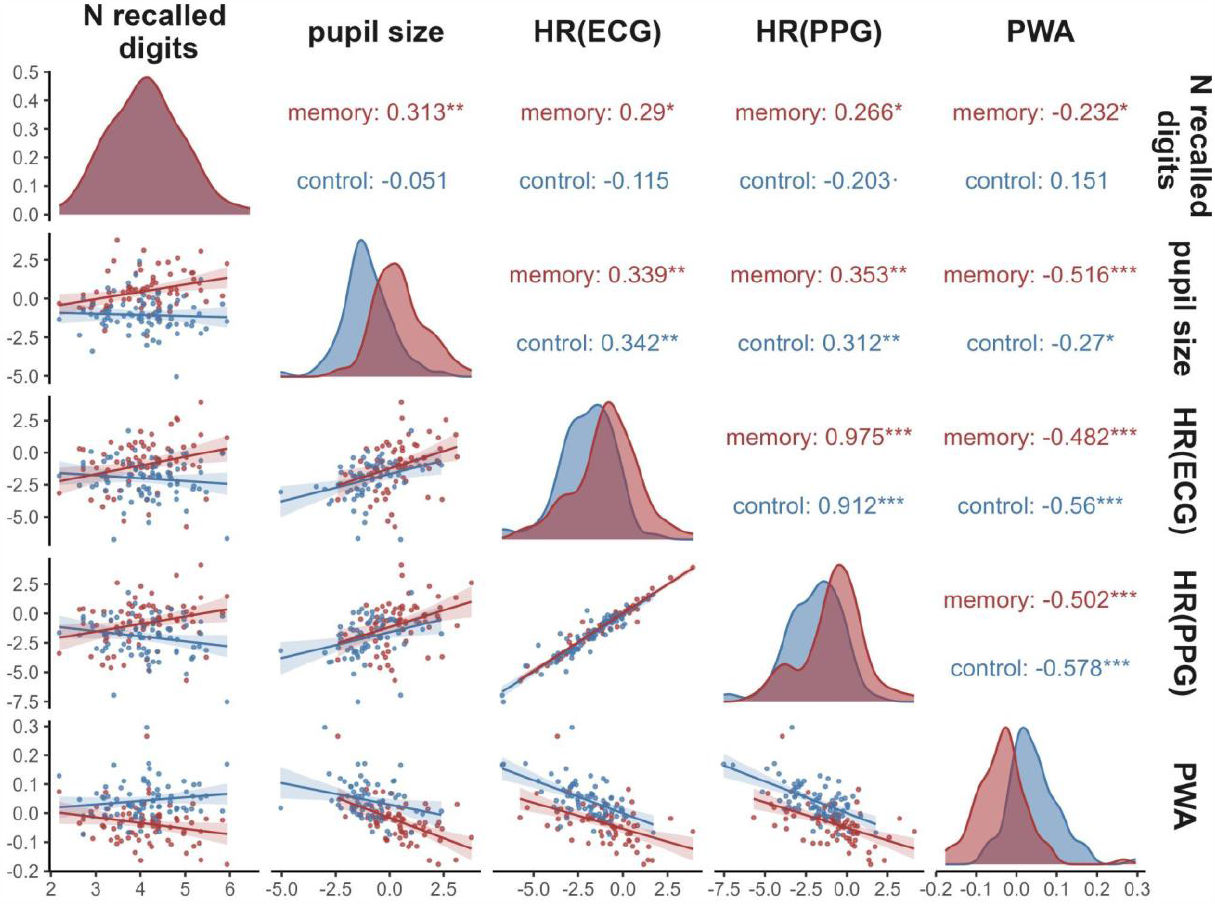
Correlations between the variables, density distributions, and scatterplots for each of the variables. The physiological parameters are averaged over all conditions and times (from presentation of the first digit till end of the retention interval after the last digit in the sequence). · p < 0.1 * p < 0.05, ** p < 0.01, *** p < 0.001

## Discussion

In this study, we aimed to assess the effectiveness of PWA as a measure for tracking cognitive load in comparison to the commonly used cardiovascular reactivity measure, HR. Unlike HR, which showed no significant changes with difficulty of the task, we observed a strong suppression of PWA with increasing memory load. Moreover, when memory load exceeded the capacity of working memory, this pattern reversed, indicating memory overload. Additionally, only PWA during the memory task correlated with self-reported cognitive load. Furthermore, both HR and PWA demonstrated a relationship with behavioral performance, as a higher task-evoked HR and lower PWA were associated with a greater number of correctly recalled memory items. These findings strongly support the notion that PWA is sensitive to cognitive overload and holds significant potential for practical application due to the ease of obtaining PPG data.

Peripheral vessel tone as expressed in PWA was changing as a function of cognitive task and load. Among all studied physiological parameters, only PWA was related to self-assessed expression of cognitive load during the tasks as measured by NASA-TLX scales. The task-evoked PWA not only decreased in response to cognitive load but also displayed a pattern of overload that was reflected in a reverse of the trajectory of PWA after about 11-digit load. This uptrend in PWA can be interpreted as a release of effort due to cognitive overload. Although the exact timing of this event may not be precisely detected by PWA, the fact that the overload was detected makes the measure an important non-invasive marker of cognitive load, largely overlooked by previous studies.

According to motivation intensity theory, effort involves the mobilization of resources for instrumental behavior and is closely linked to the activation of the sympathetic nervous system (Gendolla & Wright, 2012). The current study’s findings demonstrate that effort-mediated changes in peripheral vasoconstriction (as expressed in suppression of PWA) align with this theory, as vasoconstriction is solely regulated by sympathetic activations (Babchenko et al., 2001; Shelley, 2007). Yet, it has been suggested that effort is exclusively associated with beta-adrenergic sympathetic activation (e.g., Obrist, 1976), as supported by previous studies employing the pre-ejection period, an index of beta-adrenergic impact (e.g., Richter et al., 2008). The fact that PWA is predominantly influenced by alpha-adrenergic sympathetic activation (Guyton, 1977) appears as a potential conflict with the traditional beta-adrenergic perspective. However, previous research have similarly demonstrated systematic effects of effortful cognitive task demands on indices of peripheral vasoconstriction (Iani et al., 2004; Kemper et al., 2008; Waldstein et al., 1997). We suggest that performance in at least some cognitive tasks is supported by alpha-adrenergic activation, but future research is needed to compare the pre-ejection period and PWA within a single study to gain a comprehensive understanding of the relationship between various adrenergic impacts in cognitive tasks.

We have demonstrated the comparable effectiveness of task-evoked HR measurements using PPG and ECG, suggesting the interchangeability of these measures in mental load research. However, heart rate was not sensitive to small changes in cognitive load and associated mental effort. In studies examining cognitive load and its impact on physiological responses, it has been observed that HR is capable of distinguishing between control and memory tasks but lacks sensitivity in detecting differences between levels of cognitive load. This finding aligns with a working memory study conducted by Iani et al.(2004), which also failed to identify a general effect of memory load on HR. This lack of sensitivity may be due to the influence of different branches of ANS: the simultaneous activity of both branches of the ANS may have canceled out the weaker effects of memory load (as compared with the effect of task) on HR.

HR strongly differed between control and memory tasks independently of whether baseline normalization was applied or not. Baseline normalization appeared to be more crucial for accurate PWA measurements compared to HR. Large individual variability of indexes derived from the pulse wave has been reported previously (Lyu et al., 2015). The dramatic improvement in the assessment of cognitive load in PWA after baseline normalization can shed light on the mixed results observed in previous studies that aimed to differentiate between levels of task difficulty (Iani et al., 2004; Xuan et al., 2020). Therefore, we suggest that future investigations consider trial-level baseline PWA values for accurate comparisons across cognitive load levels and to establish consistent associations with cardiovascular responses.

In our previous study, we showed that a well-established correlate of effort and cognitive load – pupil dilation – peaked at 8 items load before a period of constriction returning pupil size to baseline level. Here, PWA peaked after the presentation of the 10th digit. At the 11^th^ and 12^th^ digit, both pupil response and PWA reversed their patterns observed with lower load levels. HR remained roughly on the same level after presentation of the fifth digit in the sequence. It is known that pupil size is influenced by both the sympathetic and parasympathetic nervous systems. Steinhauer et al. (2004) have come to the conclusion that emotional arousal primarily affects pupil dilation through sympathetic activation, while the similar dilation in response to cognitive load is primarily affected by the inhibition of the parasympathetic system. However, the current study revealed a strong correlation between task-evoked pupil size and PWA, which included, first, their simultaneous increase with cognitive load, and second, a similar drop in the amplitude of both responses in the most challenging conditions, thus indicating a shared underlying mechanism. The similarity between pupil size and PWA is additionally stressed by the fact that no parallel drop of response was observed in HR, controlled by both autonomous branches. We propose, therefore, that alpha adrenergic sympathetic activation is involved in both task-evoked pupil dilation and peripheral vasoconstriction during cognitive tasks. This activation grows with increasing load and decreases with overload.

As with all exploratory studies, ours has certain limitations. One significant limitation is that PWA can be influenced by various non-cognitive factors including physical activity, emotional stress, and sleepiness. Consequently, our findings, which were gathered during a state of “normal” wakefulness, may not apply to situations involving simultaneous cognitive and physical stressors. Additionally, the potential for generalizing our results is somewhat restricted because 88% of our study participants were female. While current evidence does not indicate significant gender-based differences in physiological responses to cognitive load, the possibility of such differences cannot be entirely dismissed.

To summarize, PWA and pupil size have emerged as valuable metrics for investigating the intensity of effort, both reflecting the response of the sympathetic nervous system to the increased cognitive load. Compared with the pupil dilation, PWA offers advantages such as simplicity, independence from luminosity and lighting conditions, and versatility. The alternative means for monitoring sympathetic activation such as heart rate variability and skin conductance operate on much larger time scales, while measuring pre-ejection period is distinctively more technologically challenging. In contrast, the PWA response is immediate, and the recording of PPG is possible with modern smartphones and, thanks to recent developments, via a web camera remotely (Djeldjli et al., 2021; van Es et al., 2023), which opens up new avenues for research and practical utilization of these measures. These developments have the potential to expand the application of PPG and PWA to enhance our understanding and assessment of cognitive load in various contexts.

## Author contributions statement

YGP: Conceptualization, Funding acquisition, Data curation, Formal analysis, Project administration, Supervision, Visualization, Methodology, Writing - original draft, Writing - review & editing.

ASG: Investigation, Formal analysis, Writing - original draft, Writing - review & editing.

AIKos, AIKot, DK: Investigation, Methodology, Data curation, Writing - review & editing.

BK: Writing - review & editing.

## Data availability statement

The data are publicly available on OpenNeuro (https://openneuro.org/datasets/ds003838).

## Acknowledgements

The research funding from the Ministry of Science and Higher Education of the Russian Federation (Ural Federal University Program of Development within the Priority-2030 Program) is gratefully acknowledged.

