## Supplementary for "Task-evoked pulse wave amplitude tracks cognitive load"

### Supplementary materials

Table S1 – Pairwise comparison of relative pulse wave amplitude in 2 second time windows after digit presentation averaged over all sequences. Note – uncorrected p-values are reported.

| Contrast (digit serial positions) | Estimate | SE | *t (77)* | *p* |
|---|---|---|---|---|
| | | **Control task** | | |
| 1 - 2 | -0.01414 | 0.003424 | -4.12953 | 9.14E-05 |
| 1 - 3 | -0.03716 | 0.005871 | -6.32941 | 1.50E-08 |
| 1 - 4 | -0.0587 | 0.0082 | -7.1582 | 4.16E-10 |
| 1 - 5 | -0.07396 | 0.009783 | -7.56018 | 7.12E-11 |
| 1 - 6 | -0.08309 | 0.010704 | -7.76312 | 2.90E-11 |
| 1 - 7 | -0.09147 | 0.011376 | -8.04085 | 8.49E-12 |
| 1 - 8 | -0.09359 | 0.011334 | -8.25756 | 3.24E-12 |
| 1 - 9 | -0.0902 | 0.011654 | -7.74006 | 3.22E-11 |
| 1 - 10 | -0.07859 | 0.012787 | -6.14637 | 3.26E-08 |
| 1 - 11 | -0.07936 | 0.012164 | -6.52438 | 6.51E-09 |
| 1 - 12 | -0.07301 | 0.012294 | -5.939 | 7.80E-08 |
| 1 - 13 | -0.06963 | 0.012559 | -5.54452 | 3.98E-07 |
| 2 - 3 | -0.02302 | 0.003215 | -7.16065 | 4.12E-10 |
| 2 - 4 | -0.04456 | 0.005983 | -7.44706 | 1.17E-10 |
| 2 - 5 | -0.05982 | 0.007858 | -7.61277 | 5.64E-11 |
| 2 - 6 | -0.06895 | 0.009179 | -7.5122 | 8.79E-11 |
| 2 - 7 | -0.07733 | 0.010046 | -7.69718 | 3.89E-11 |
| 2 - 8 | -0.07945 | 0.010337 | -7.68585 | 4.09E-11 |
| 2 - 9 | -0.07606 | 0.010643 | -7.14673 | 4.38E-10 |
| 2 - 10 | -0.06445 | 0.011765 | -5.47817 | 5.21E-07 |
| 2 - 11 | -0.06522 | 0.01138 | -5.73091 | 1.85E-07 |
| 2 - 12 | -0.05887 | 0.01154 | -5.10134 | 2.36E-06 |
| 2 - 13 | -0.05549 | 0.011809 | -4.69895 | 1.12E-05 |
| 3 - 4 | -0.02154 | 0.003263 | -6.60121 | 4.68E-09 |
| 3 - 5 | -0.0368 | 0.005511 | -6.67874 | 3.35E-09 |
| 3 - 6 | -0.04593 | 0.007394 | -6.21255 | 2.46E-08 |
| 3 - 7 | -0.05431 | 0.008299 | -6.54443 | 5.98E-09 |
| 3 - 8 | -0.05643 | 0.00872 | -6.47142 | 8.18E-09 |
| 3 - 9 | -0.05304 | 0.009077 | -5.84407 | 1.16E-07 |
| 3 - 10 | -0.04143 | 0.010506 | -3.94393 | 0.000175 |
| 3 - 11 | -0.0422 | 0.01038 | -4.06556 | 0.000115 |
| 3 - 12 | -0.03585 | 0.010473 | -3.42313 | 0.000995 |
| 3 - 13 | -0.03247 | 0.010831 | -2.99815 | 0.003656 |

| | | | | |
|---|---|---|---|---|
| 4 - 5 | -0.01526 | 0.002854 | -5.34891 | 8.80E-07 |
| 4 - 6 | -0.0244 | 0.005419 | -4.50159 | 2.36E-05 |
| 4 - 7 | -0.03277 | 0.006294 | -5.20658 | 1.55E-06 |
| 4 - 8 | -0.03489 | 0.006888 | -5.06516 | 2.72E-06 |
| 4 - 9 | -0.03151 | 0.007304 | -4.31374 | 4.71E-05 |
| 4 - 10 | -0.01989 | 0.009414 | -2.1132 | 0.037824 |
| 4 - 11 | -0.02066 | 0.009563 | -2.16056 | 0.03384 |
| 4 - 12 | -0.01431 | 0.009666 | -1.48072 | 0.142763 |
| 4 - 13 | -0.01093 | 0.010148 | -1.07734 | 0.284694 |
| 5 - 6 | -0.00913 | 0.00426 | -2.14337 | 0.035241 |
| 5 - 7 | -0.01751 | 0.005088 | -3.44059 | 0.000941 |
| 5 - 8 | -0.01963 | 0.005868 | -3.34424 | 0.001277 |
| 5 - 9 | -0.01624 | 0.00631 | -2.57367 | 0.011983 |
| 5 - 10 | -0.00463 | 0.008731 | -0.53025 | 0.597466 |
| 5 - 11 | -0.0054 | 0.009005 | -0.59933 | 0.550712 |
| 5 - 12 | 0.000952 | 0.009134 | 0.104278 | 0.91722 |
| 5 - 13 | 0.004331 | 0.009559 | 0.453078 | 0.651767 |
| 6 - 7 | -0.00837 | 0.002368 | -3.53685 | 0.00069 |
| 6 - 8 | -0.01049 | 0.00391 | -2.68402 | 0.008902 |
| 6 - 9 | -0.00711 | 0.004994 | -1.42373 | 0.158563 |
| 6 - 10 | 0.004502 | 0.00828 | 0.543701 | 0.588218 |
| 6 - 11 | 0.003734 | 0.008678 | 0.430329 | 0.668159 |
| 6 - 12 | 0.010084 | 0.008749 | 1.152554 | 0.252659 |
| 6 - 13 | 0.013462 | 0.009121 | 1.475979 | 0.144027 |
| 7 - 8 | -0.00212 | 0.002314 | -0.91564 | 0.362717 |
| 7 - 9 | 0.001265 | 0.003688 | 0.34302 | 0.732518 |
| 7 - 10 | 0.012877 | 0.007707 | 1.670751 | 0.098831 |
| 7 - 11 | 0.012109 | 0.008192 | 1.478085 | 0.143464 |
| 7 - 12 | 0.018458 | 0.008447 | 2.185167 | 0.031918 |
| 7 - 13 | 0.021837 | 0.008959 | 2.437331 | 0.017103 |
| 8 - 9 | 0.003384 | 0.002423 | 1.396921 | 0.16645 |
| 8 - 10 | 0.014996 | 0.007279 | 2.060088 | 0.042769 |
| 8 - 11 | 0.014228 | 0.00777 | 1.831201 | 0.070939 |
| 8 - 12 | 0.020578 | 0.008041 | 2.558982 | 0.012459 |
| 8 - 13 | 0.023956 | 0.008549 | 2.80207 | 0.00642 |
| 9 - 10 | 0.011612 | 0.006691 | 1.735379 | 0.086674 |
| 9 - 11 | 0.010844 | 0.007229 | 1.500026 | 0.137697 |
| 9 - 12 | 0.017193 | 0.007703 | 2.232064 | 0.028518 |
| 9 - 13 | 0.020572 | 0.008455 | 2.433242 | 0.017283 |
| 10 - 11 | -0.00077 | 0.00284 | -0.27032 | 0.787638 |
| 10 - 12 | 0.005582 | 0.004599 | 1.213821 | 0.228526 |
| 10 - 13 | 0.00896 | 0.006416 | 1.396573 | 0.166555 |
| 11 - 12 | 0.006349 | 0.003448 | 1.841399 | 0.069414 |
| 11 - 13 | 0.009728 | 0.005652 | 1.721252 | 0.089221 |
| 12 - 13 | 0.003379 | 0.00326 | 1.036427 | 0.303247 |

| Memory task | | | | |
|---|---|---|---|---|
| 1 - 2 | 0.003119 | 0.003019 | 1.03318 | 0.304754 |
| 1 - 3 | 0.000858 | 0.004873 | 0.176022 | 0.860739 |
| 1 - 4 | -0.00219 | 0.006323 | -0.34584 | 0.730405 |
| 1 - 5 | 0.003589 | 0.007595 | 0.472478 | 0.637922 |
| 1 - 6 | 0.017439 | 0.009476 | 1.840337 | 0.069572 |
| 1 - 7 | 0.042186 | 0.010607 | 3.977105 | 0.000156 |
| 1 - 8 | 0.063826 | 0.011851 | 5.385565 | 7.59E-07 |
| 1 - 9 | 0.081418 | 0.012635 | 6.443963 | 9.20E-09 |
| 1 - 10 | 0.091695 | 0.013305 | 6.891786 | 1.33E-09 |
| 1 - 11 | 0.091649 | 0.013735 | 6.672689 | 3.44E-09 |
| 1 - 12 | 0.084093 | 0.013811 | 6.088995 | 4.15E-08 |
| 1 - 13 | 0.074938 | 0.014273 | 5.250226 | 1.31E-06 |
| 2 - 3 | -0.00226 | 0.002403 | -0.94083 | 0.349732 |
| 2 - 4 | -0.00531 | 0.004189 | -1.26645 | 0.20917 |
| 2 - 5 | 0.00047 | 0.005848 | 0.08033 | 0.936183 |
| 2 - 6 | 0.01432 | 0.008156 | 1.755721 | 0.083113 |
| 2 - 7 | 0.039067 | 0.009686 | 4.033295 | 0.000128 |
| 2 - 8 | 0.060708 | 0.011228 | 5.406758 | 6.96E-07 |
| 2 - 9 | 0.078299 | 0.012225 | 6.404589 | 1.09E-08 |
| 2 - 10 | 0.088576 | 0.013069 | 6.777343 | 2.19E-09 |
| 2 - 11 | 0.08853 | 0.013607 | 6.506407 | 7.04E-09 |
| 2 - 12 | 0.080974 | 0.013752 | 5.888018 | 9.65E-08 |
| 2 - 13 | 0.071819 | 0.014079 | 5.101211 | 2.36E-06 |
| 3 - 4 | -0.00304 | 0.002155 | -1.41291 | 0.161712 |
| 3 - 5 | 0.002731 | 0.004097 | 0.666509 | 0.507078 |
| 3 - 6 | 0.016581 | 0.00682 | 2.431144 | 0.017376 |
| 3 - 7 | 0.041328 | 0.008685 | 4.758696 | 8.94E-06 |
| 3 - 8 | 0.062969 | 0.010466 | 6.016559 | 5.63E-08 |
| 3 - 9 | 0.08056 | 0.011684 | 6.894965 | 1.31E-09 |
| 3 - 10 | 0.090838 | 0.012663 | 7.173437 | 3.90E-10 |
| 3 - 11 | 0.090792 | 0.013326 | 6.813338 | 1.87E-09 |
| 3 - 12 | 0.083236 | 0.013482 | 6.173906 | 2.90E-08 |
| 3 - 13 | 0.07408 | 0.013726 | 5.396986 | 7.25E-07 |
| 4 - 5 | 0.005775 | 0.002413 | 2.393918 | 0.019103 |
| 4 - 6 | 0.019626 | 0.005426 | 3.61702 | 0.00053 |
| 4 - 7 | 0.044372 | 0.007503 | 5.913949 | 8.66E-08 |
| 4 - 8 | 0.066013 | 0.009531 | 6.925854 | 1.15E-09 |
| 4 - 9 | 0.083605 | 0.010985 | 7.610785 | 5.69E-11 |
| 4 - 10 | 0.093882 | 0.012122 | 7.744896 | 3.15E-11 |
| 4 - 11 | 0.093836 | 0.012915 | 7.265635 | 2.60E-10 |
| 4 - 12 | 0.08628 | 0.013091 | 6.590855 | 4.90E-09 |
| 4 - 13 | 0.077124 | 0.013315 | 5.79228 | 1.44E-07 |
| 5 - 6 | 0.01385 | 0.003522 | 3.932172 | 0.000183 |
| 5 - 7 | 0.038597 | 0.005764 | 6.695707 | 3.11E-09 |

| | | | | |
|---|---|---|---|---|
| 5 - 8 | 0.060238 | 0.008005 | 7.52494 | 8.31E-11 |
| 5 - 9 | 0.077829 | 0.009741 | 7.989646 | 1.06E-11 |
| 5 - 10 | 0.088107 | 0.011045 | 7.976923 | 1.13E-11 |
| 5 - 11 | 0.088061 | 0.011974 | 7.354611 | 1.76E-10 |
| 5 - 12 | 0.080505 | 0.012241 | 6.576458 | 5.21E-09 |
| 5 - 13 | 0.071349 | 0.01247 | 5.721777 | 1.92E-07 |
| 6 - 7 | 0.024747 | 0.002931 | 8.443765 | 1.42E-12 |
| 6 - 8 | 0.046388 | 0.005732 | 8.093321 | 6.72E-12 |
| 6 - 9 | 0.063979 | 0.007876 | 8.122845 | 5.90E-12 |
| 6 - 10 | 0.074256 | 0.009559 | 7.768439 | 2.84E-11 |
| 6 - 11 | 0.07421 | 0.010729 | 6.916639 | 1.20E-09 |
| 6 - 12 | 0.066654 | 0.011149 | 5.978322 | 6.61E-08 |
| 6 - 13 | 0.057499 | 0.01145 | 5.021656 | 3.23E-06 |
| 7 - 8 | 0.021641 | 0.003314 | 6.529452 | 6.37E-09 |
| 7 - 9 | 0.039233 | 0.005759 | 6.812553 | 1.88E-09 |
| 7 - 10 | 0.04951 | 0.007855 | 6.302665 | 1.68E-08 |
| 7 - 11 | 0.049464 | 0.009178 | 5.389291 | 7.47E-07 |
| 7 - 12 | 0.041908 | 0.009754 | 4.296396 | 5.02E-05 |
| 7 - 13 | 0.032752 | 0.010251 | 3.195111 | 0.002027 |
| 8 - 9 | 0.017592 | 0.002896 | 6.07542 | 4.40E-08 |
| 8 - 10 | 0.027869 | 0.005664 | 4.92043 | 4.79E-06 |
| 8 - 11 | 0.027823 | 0.007056 | 3.943336 | 0.000176 |
| 8 - 12 | 0.020267 | 0.007831 | 2.587947 | 0.011536 |
| 8 - 13 | 0.011111 | 0.008604 | 1.29135 | 0.200446 |
| 9 - 10 | 0.010277 | 0.004049 | 2.538227 | 0.01316 |
| 9 - 11 | 0.010231 | 0.005312 | 1.926177 | 0.057773 |
| 9 - 12 | 0.002675 | 0.006289 | 0.425373 | 0.671752 |
| 9 - 13 | -0.00648 | 0.007249 | -0.89399 | 0.374111 |
| 10 - 11 | -4.61E-05 | 0.002315 | -0.01989 | 0.984181 |
| 10 - 12 | -0.0076 | 0.003886 | -1.95619 | 0.054069 |
| 10 - 13 | -0.01676 | 0.005321 | -3.14942 | 0.002329 |
| 11 - 12 | -0.00756 | 0.002341 | -3.22803 | 0.001832 |
| 11 - 13 | -0.01671 | 0.004065 | -4.11067 | 9.78E-05 |
| 12 - 13 | -0.00916 | 0.00251 | -3.64833 | 0.000478 |

Table S2 – Pairwise comparison of relative heart rate in 2 second time windows after digit presentation averaged over all sequences. Note – uncorrected p-values are reported.

| Contrast (digit serial positions) | Estimate | SE | *t (77)* | *p* |
|---|---|---|---|---|
| | | **Control task** | | |
| 1 - 2 | 0.539574 | 0.103127 | 5.232144 | 1.40E-06 |
| 1 - 3 | 1.115654 | 0.137747 | 8.099311 | 6.55E-12 |
| 1 - 4 | 1.462867 | 0.154542 | 9.465799 | 1.52E-14 |
| 1 - 5 | 1.830278 | 0.14695 | 12.45507 | 3.90E-20 |
| 1 - 6 | 2.159818 | 0.166749 | 12.95248 | 5.01E-21 |
| 1 - 7 | 2.427863 | 0.182738 | 13.28607 | 1.29E-21 |
| 1 - 8 | 2.497191 | 0.171465 | 14.56381 | 7.98E-24 |
| 1 - 9 | 2.630301 | 0.178426 | 14.74173 | 4.00E-24 |
| 1 - 10 | 2.345458 | 0.213265 | 10.99788 | 1.88E-17 |
| 1 - 11 | 2.56682 | 0.230934 | 11.11496 | 1.13E-17 |
| 1 - 12 | 2.75577 | 0.227824 | 12.09606 | 1.75E-19 |
| 1 - 13 | 2.687528 | 0.227044 | 11.83703 | 5.21E-19 |
| 2 - 3 | 0.57608 | 0.094485 | 6.097074 | 4.02E-08 |
| 2 - 4 | 0.923293 | 0.125251 | 7.371539 | 1.63E-10 |
| 2 - 5 | 1.290703 | 0.130291 | 9.906337 | 2.18E-15 |
| 2 - 6 | 1.620244 | 0.150572 | 10.76062 | 5.23E-17 |
| 2 - 7 | 1.888289 | 0.174106 | 10.84562 | 3.62E-17 |
| 2 - 8 | 1.957617 | 0.157567 | 12.42399 | 4.44E-20 |
| 2 - 9 | 2.090727 | 0.169117 | 12.3626 | 5.74E-20 |
| 2 - 10 | 1.805883 | 0.195767 | 9.224671 | 4.43E-14 |
| 2 - 11 | 2.027245 | 0.223527 | 9.069337 | 8.82E-14 |
| 2 - 12 | 2.216196 | 0.218513 | 10.14218 | 7.75E-16 |
| 2 - 13 | 2.147954 | 0.221425 | 9.7006 | 5.40E-15 |
| 3 - 4 | 0.347213 | 0.07888 | 4.401794 | 3.41E-05 |
| 3 - 5 | 0.714623 | 0.105644 | 6.764438 | 2.31E-09 |
| 3 - 6 | 1.044164 | 0.136087 | 7.672772 | 4.33E-11 |
| 3 - 7 | 1.312209 | 0.14368 | 9.132871 | 6.65E-14 |
| 3 - 8 | 1.381537 | 0.130088 | 10.62002 | 9.63E-17 |
| 3 - 9 | 1.514647 | 0.14315 | 10.58083 | 1.14E-16 |
| 3 - 10 | 1.229803 | 0.185752 | 6.620672 | 4.31E-09 |
| 3 - 11 | 1.451165 | 0.201284 | 7.209543 | 3.33E-10 |
| 3 - 12 | 1.640116 | 0.198026 | 8.282321 | 2.91E-12 |
| 3 - 13 | 1.571874 | 0.20844 | 7.541139 | 7.74E-11 |
| 4 - 5 | 0.367411 | 0.071011 | 5.173983 | 1.77E-06 |
| 4 - 6 | 0.696952 | 0.110282 | 6.319725 | 1.56E-08 |
| 4 - 7 | 0.964997 | 0.12452 | 7.749728 | 3.08E-11 |
| 4 - 8 | 1.034324 | 0.118935 | 8.696513 | 4.61E-13 |

| | | | | |
|---|---|---|---|---|
| 4 - 9 | 1.167434 | 0.124745 | 9.358583 | 2.45E-14 |
| 4 - 10 | 0.882591 | 0.175868 | 5.018497 | 3.27E-06 |
| 4 - 11 | 1.103953 | 0.19152 | 5.764178 | 1.61E-07 |
| 4 - 12 | 1.292904 | 0.193908 | 6.667619 | 3.52E-09 |
| 4 - 13 | 1.224662 | 0.194484 | 6.296964 | 1.72E-08 |
| 5 - 6 | 0.329541 | 0.101191 | 3.256622 | 0.001678 |
| 5 - 7 | 0.597586 | 0.116003 | 5.151453 | 1.94E-06 |
| 5 - 8 | 0.666913 | 0.124826 | 5.342758 | 9.02E-07 |
| 5 - 9 | 0.800024 | 0.115834 | 6.906615 | 1.25E-09 |
| 5 - 10 | 0.51518 | 0.173479 | 2.969697 | 0.003973 |
| 5 - 11 | 0.736542 | 0.188183 | 3.913963 | 0.000195 |
| 5 - 12 | 0.925493 | 0.181822 | 5.0901 | 2.47E-06 |
| 5 - 13 | 0.857251 | 0.185134 | 4.63043 | 1.46E-05 |
| 6 - 7 | 0.268045 | 0.080029 | 3.349367 | 0.001256 |
| 6 - 8 | 0.337372 | 0.101671 | 3.318268 | 0.001385 |
| 6 - 9 | 0.470483 | 0.099305 | 4.737746 | 9.68E-06 |
| 6 - 10 | 0.185639 | 0.160745 | 1.154869 | 0.251716 |
| 6 - 11 | 0.407001 | 0.17486 | 2.327577 | 0.022563 |
| 6 - 12 | 0.595952 | 0.181921 | 3.275886 | 0.001581 |
| 6 - 13 | 0.52771 | 0.189252 | 2.788394 | 0.006671 |
| 7 - 8 | 0.069327 | 0.092595 | 0.748717 | 0.456309 |
| 7 - 9 | 0.202438 | 0.092871 | 2.179766 | 0.032331 |
| 7 - 10 | -0.08241 | 0.175393 | -0.46984 | 0.639801 |
| 7 - 11 | 0.138956 | 0.169984 | 0.817467 | 0.416184 |
| 7 - 12 | 0.327907 | 0.174897 | 1.874857 | 0.064604 |
| 7 - 13 | 0.259665 | 0.184969 | 1.403831 | 0.164389 |
| 8 - 9 | 0.13311 | 0.08082 | 1.646991 | 0.103635 |
| 8 - 10 | -0.15173 | 0.16623 | -0.91279 | 0.364202 |
| 8 - 11 | 0.069629 | 0.174792 | 0.398351 | 0.691474 |
| 8 - 12 | 0.25858 | 0.171395 | 1.508673 | 0.135475 |
| 8 - 13 | 0.190338 | 0.176439 | 1.078775 | 0.284056 |
| 9 - 10 | -0.28484 | 0.157107 | -1.81305 | 0.073721 |
| 9 - 11 | -0.06348 | 0.171804 | -0.3695 | 0.712768 |
| 9 - 12 | 0.125469 | 0.169282 | 0.741183 | 0.460837 |
| 9 - 13 | 0.057227 | 0.173383 | 0.330062 | 0.742249 |
| 10 - 11 | 0.221362 | 0.124016 | 1.784949 | 0.078208 |
| 10 - 12 | 0.410313 | 0.147589 | 2.780096 | 0.006827 |
| 10 - 13 | 0.342071 | 0.145359 | 2.353279 | 0.021162 |
| 11 - 12 | 0.188951 | 0.108531 | 1.740984 | 0.085681 |
| 11 - 13 | 0.120709 | 0.133864 | 0.901725 | 0.370015 |
| 12 - 13 | -0.06824 | 0.107263 | -0.63621 | 0.526524 |
| **Memory task** | | | | |
| 1 - 2 | 0.44421 | 0.111909 | 3.969375 | 0.000161 |
| 1 - 3 | 0.65613 | 0.159732 | 4.107684 | 9.88E-05 |
| 1 - 4 | 0.495847 | 0.207687 | 2.387479 | 0.019417 |

| | | | | |
|---|---|---|---|---|
| 1 - 5 | 0.354768 | 0.210255 | 1.687326 | 0.095588 |
| 1 - 6 | 0.343891 | 0.238281 | 1.443216 | 0.153016 |
| 1 - 7 | 0.350501 | 0.254932 | 1.374883 | 0.173155 |
| 1 - 8 | 0.187088 | 0.274613 | 0.681276 | 0.497741 |
| 1 - 9 | 0.154395 | 0.293222 | 0.526545 | 0.600023 |
| 1 - 10 | 0.124237 | 0.305564 | 0.406583 | 0.685442 |
| 1 - 11 | 0.284701 | 0.307888 | 0.924688 | 0.358017 |
| 1 - 12 | 0.396767 | 0.327967 | 1.209776 | 0.230066 |
| 1 - 13 | 0.408692 | 0.335531 | 1.218046 | 0.226926 |
| 2 - 3 | 0.21192 | 0.085754 | 2.471263 | 0.015673 |
| 2 - 4 | 0.051637 | 0.15469 | 0.333812 | 0.739428 |
| 2 - 5 | -0.08944 | 0.170374 | -0.52497 | 0.60111 |
| 2 - 6 | -0.10032 | 0.199254 | -0.50347 | 0.616069 |
| 2 - 7 | -0.09371 | 0.210958 | -0.44421 | 0.658139 |
| 2 - 8 | -0.25712 | 0.232752 | -1.1047 | 0.272728 |
| 2 - 9 | -0.28982 | 0.256664 | -1.12917 | 0.262334 |
| 2 - 10 | -0.31997 | 0.280433 | -1.141 | 0.257407 |
| 2 - 11 | -0.15951 | 0.283046 | -0.56355 | 0.5747 |
| 2 - 12 | -0.04744 | 0.294339 | -0.16119 | 0.872369 |
| 2 - 13 | -0.03552 | 0.305126 | -0.1164 | 0.907635 |
| 3 - 4 | -0.16028 | 0.093116 | -1.72132 | 0.089208 |
| 3 - 5 | -0.30136 | 0.118293 | -2.54758 | 0.01284 |
| 3 - 6 | -0.31224 | 0.155492 | -2.00807 | 0.048143 |
| 3 - 7 | -0.30563 | 0.166478 | -1.83585 | 0.07024 |
| 3 - 8 | -0.46904 | 0.190313 | -2.46459 | 0.015945 |
| 3 - 9 | -0.50174 | 0.214777 | -2.33608 | 0.022091 |
| 3 - 10 | -0.53189 | 0.244851 | -2.17231 | 0.032909 |
| 3 - 11 | -0.37143 | 0.244347 | -1.52009 | 0.132585 |
| 3 - 12 | -0.25936 | 0.258018 | -1.00522 | 0.317941 |
| 3 - 13 | -0.24744 | 0.272466 | -0.90814 | 0.366636 |
| 4 - 5 | -0.14108 | 0.079847 | -1.76686 | 0.081214 |
| 4 - 6 | -0.15196 | 0.128159 | -1.18569 | 0.239391 |
| 4 - 7 | -0.14535 | 0.147851 | -0.98306 | 0.328659 |
| 4 - 8 | -0.30876 | 0.164382 | -1.8783 | 0.064125 |
| 4 - 9 | -0.34145 | 0.192811 | -1.77092 | 0.080531 |
| 4 - 10 | -0.37161 | 0.227817 | -1.63118 | 0.106936 |
| 4 - 11 | -0.21115 | 0.232867 | -0.90673 | 0.367379 |
| 4 - 12 | -0.09908 | 0.252989 | -0.39164 | 0.696407 |
| 4 - 13 | -0.08716 | 0.268207 | -0.32496 | 0.746096 |
| 5 - 6 | -0.01088 | 0.089372 | -0.12171 | 0.903448 |
| 5 - 7 | -0.00427 | 0.114235 | -0.03735 | 0.970299 |
| 5 - 8 | -0.16768 | 0.134587 | -1.2459 | 0.21658 |
| 5 - 9 | -0.20037 | 0.160654 | -1.24724 | 0.216091 |
| 5 - 10 | -0.23053 | 0.198968 | -1.15864 | 0.250185 |
| 5 - 11 | -0.07007 | 0.201314 | -0.34805 | 0.72875 |

| | | | | |
|---|---|---|---|---|
| 5 - 12 | 0.041998 | 0.227425 | 0.184669 | 0.853974 |
| 5 - 13 | 0.053924 | 0.241223 | 0.223542 | 0.823706 |
| 6 - 7 | 0.00661 | 0.076462 | 0.086449 | 0.931334 |
| 6 - 8 | -0.1568 | 0.105268 | -1.48957 | 0.140423 |
| 6 - 9 | -0.1895 | 0.132622 | -1.42884 | 0.157093 |
| 6 - 10 | -0.21965 | 0.173321 | -1.26733 | 0.208858 |
| 6 - 11 | -0.05919 | 0.170698 | -0.34676 | 0.729721 |
| 6 - 12 | 0.052876 | 0.196529 | 0.269048 | 0.788612 |
| 6 - 13 | 0.064801 | 0.216769 | 0.298939 | 0.765792 |
| 7 - 8 | -0.16341 | 0.060813 | -2.68716 | 0.008826 |
| 7 - 9 | -0.19611 | 0.093596 | -2.09525 | 0.039437 |
| 7 - 10 | -0.22626 | 0.148128 | -1.52749 | 0.130737 |
| 7 - 11 | -0.0658 | 0.146877 | -0.448 | 0.655412 |
| 7 - 12 | 0.046266 | 0.170961 | 0.270621 | 0.787406 |
| 7 - 13 | 0.058191 | 0.186085 | 0.312711 | 0.755346 |
| 8 - 9 | -0.03269 | 0.064746 | -0.50494 | 0.615041 |
| 8 - 10 | -0.06285 | 0.133528 | -0.47069 | 0.639193 |
| 8 - 11 | 0.097613 | 0.143321 | 0.681081 | 0.497864 |
| 8 - 12 | 0.209679 | 0.163433 | 1.282968 | 0.203352 |
| 8 - 13 | 0.221604 | 0.173469 | 1.277488 | 0.205268 |
| 9 - 10 | -0.03016 | 0.105372 | -0.2862 | 0.775494 |
| 9 - 11 | 0.130306 | 0.121865 | 1.069269 | 0.288289 |
| 9 - 12 | 0.242372 | 0.147915 | 1.638587 | 0.105379 |
| 9 - 13 | 0.254297 | 0.149064 | 1.705966 | 0.092045 |
| 10 - 11 | 0.160464 | 0.092211 | 1.74017 | 0.085824 |
| 10 - 12 | 0.27253 | 0.126768 | 2.149828 | 0.034708 |
| 10 - 13 | 0.284455 | 0.128818 | 2.208186 | 0.030207 |
| 11 - 12 | 0.112066 | 0.087134 | 1.286131 | 0.202251 |
| 11 - 13 | 0.123991 | 0.111903 | 1.10803 | 0.271298 |
| 12 - 13 | 0.011925 | 0.074703 | 0.159634 | 0.873587 |

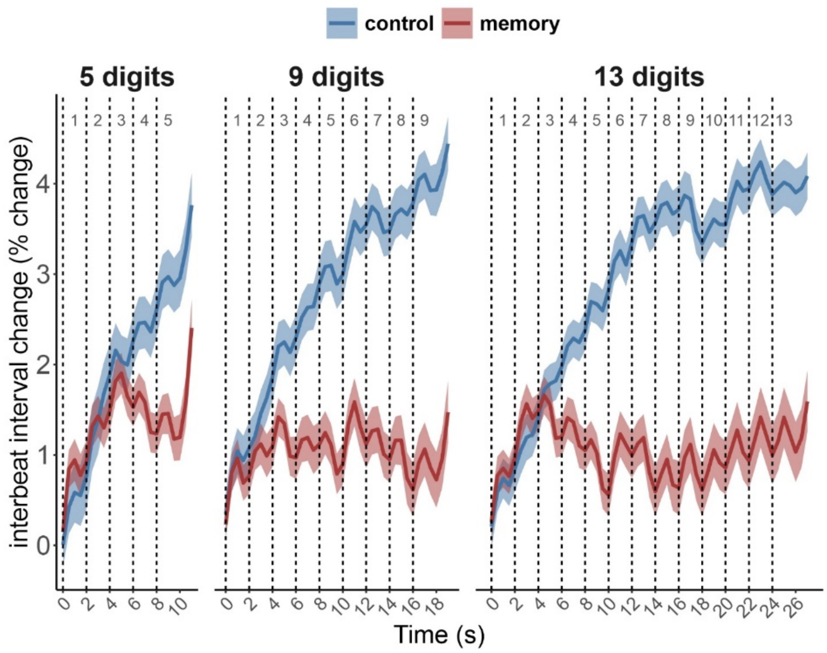


Figure S1 – Temporal dynamics of relative interbeat interval registered by ECG. Each dashed line represents the onset of a digit presentation. Time 0 indicates presentation of the first digit in the sequence.

Table S3 – The Task (control vs memory) and Load (13 levels) effects on relative interbeat intervals

| | *Interbeat interval* | | | |
|---|---|---|---|---|
| | *df* | *F* | *p* | $\eta^2$ |
| Task | 1, 77 | 46.23 | <.001 | .38 |
| Load | 2.6, 199.98 | 19.74 | <.001 | . 20 |
| Task x Load | 3.4, 261.96 | 31.55 | <.001 | . 29 |

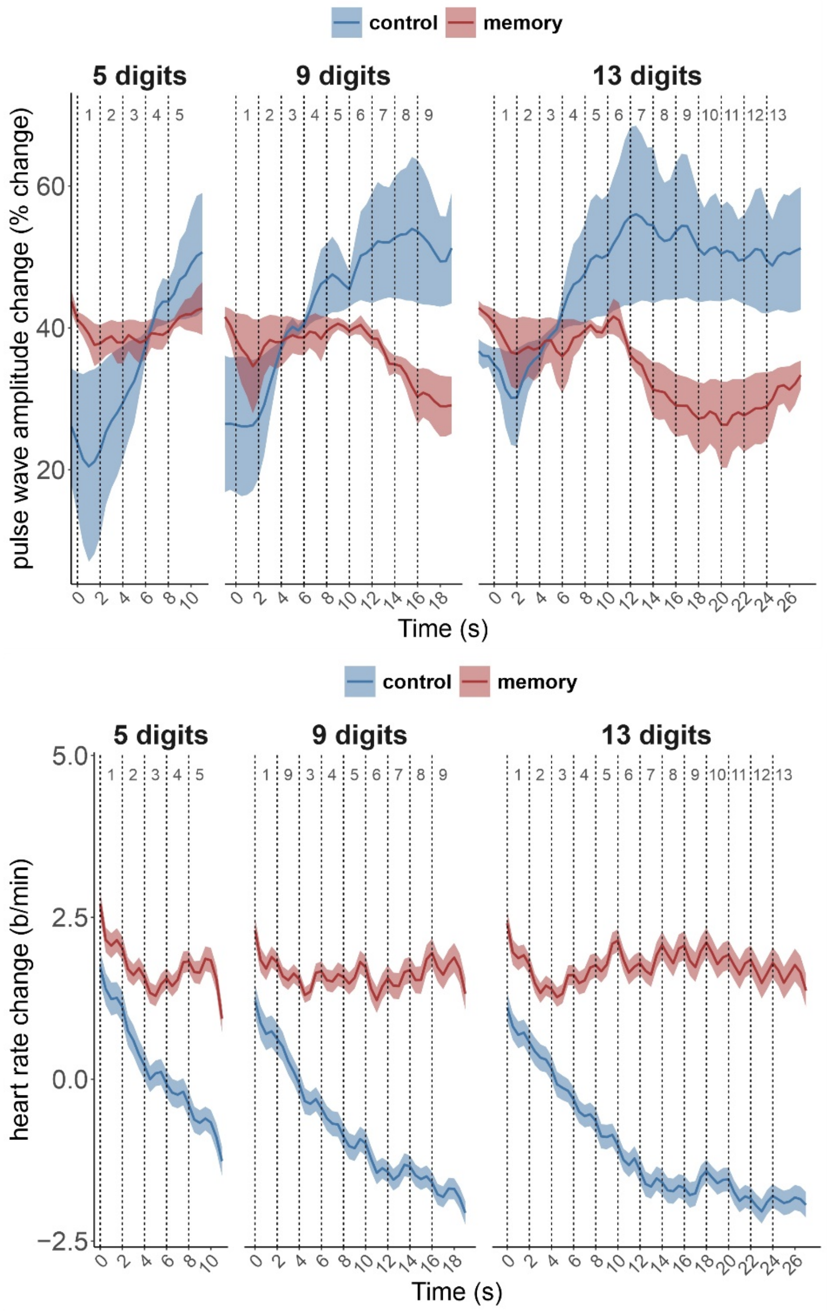


Figure S2 – Temporal dynamics of relative pulse wave amplitude (top panel) and heart rate (bottom panel) with resting state values used for baseline normalization. Each dashed line represents the onset of a digit presentation. Time 0 indicates presentation of the first digit in the sequence.

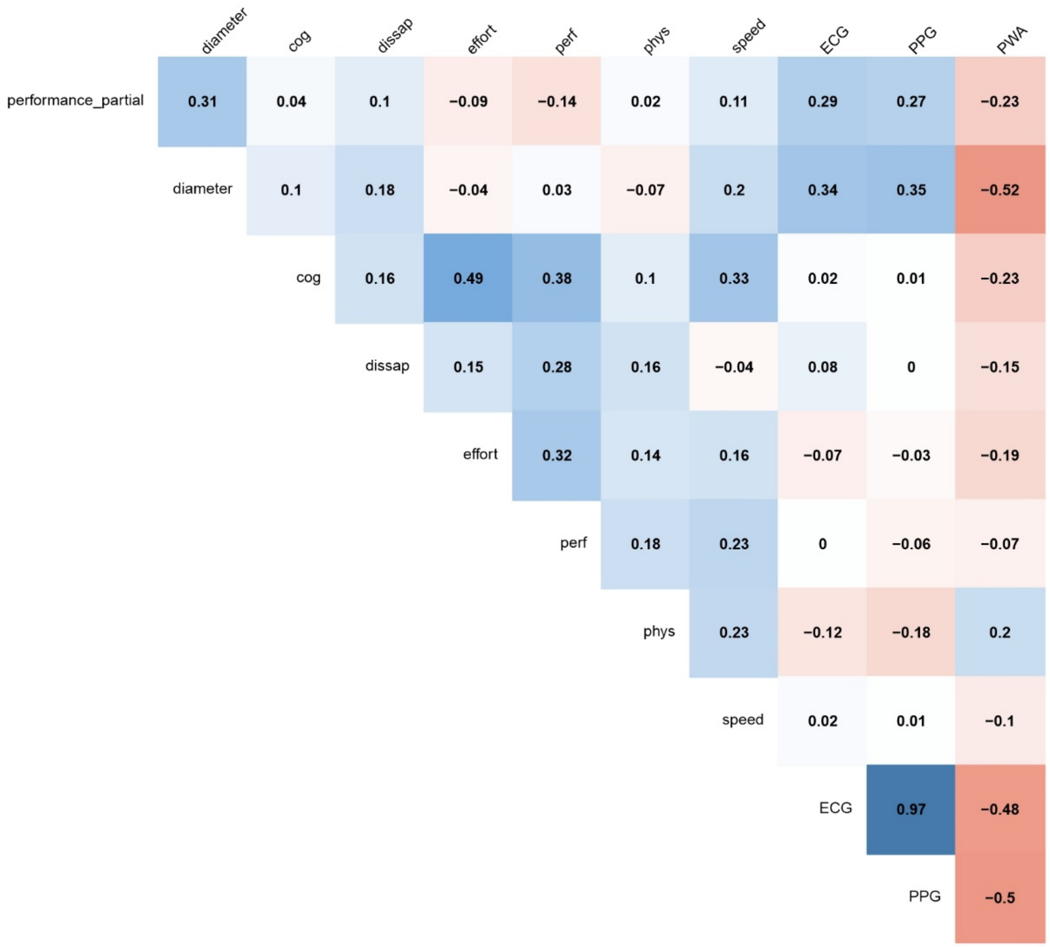


Figure S3 – Pearson correlations between physiological variables in the memory task: heart rate extracted from ECG – ECG, heart rate extracted from PPG – PPG, pulse wave amplitude – PWA, pupil size – diameter, behavioral performance (number of correctly recalled digits) – performance_partial, and NASA-TLX subscales (assessing how mentally demanding – cog, (2) frustrating – dissap, (3) effortful – effort, (4) difficult to perform – perf, (5) physically demanding – phys, and (6) temporally demanding – speed – the task was)
